# High Throughput PET/CT Imaging Using a Multiple Mouse Imaging System

**DOI:** 10.1101/602391

**Authors:** Hannah E. Greenwood, Zoltan Nyitrai, Gabor Mocsai, Sandor Hobor, Timothy H. Witney

**Author notes:** **Corresponding author:** Timothy H. Witney, Department of Imaging Chemistry and Biology, School of Biomedical Engineering and & Imaging Sciences, King’s College London, London, UK.

## Abstract

A considerable limitation of current small animal positron emission tomography/computed tomography (PET/CT) imaging is the low throughput of image acquisitions. Subsequently, to design sufficiently-powered studies, high costs accumulate. Together with Mediso Medical Imaging Systems, a four-bed mouse ‘hotel’ was developed to simultaneously image up to four mice, thereby reducing the cost and maximising radiotracer usage when compared to scans performed with a single mouse bed.

**Methods:** For physiological evaluation of the four-bed mouse hotel, temperature and anaesthesia were tested for uniformity, followed by [^18^F]fluorodeoxyglucose (FDG) PET/CT imaging of ‘mini’ image quality (IQ) phantoms specifically designed to fit the new imaging system. Post-reconstruction, National Electrical Manufacturers Association (NEMA) NU-4 tests examined uniformity, recovery coefficients (RCs) and spill-over ratios (SORs). To evaluate the bed under standard *in vivo* imaging conditions, four mice were simultaneously scanned by dynamic [^18^F]FDG PET/CT over 60 minutes using the four-bed mouse hotel, with quantified images compared to those acquired using a single mouse bed.

**Results:** The bed maintained a constant temperature of 36.8°C ± 0.4°C (*n* = 4), with anaesthesia distributed evenly to each nose cone (2.9 ± 0.1 L/min, *n* = 4). The NEMA tests performed on reconstructed mini IQ phantom images acquired using the four-bed mouse hotel revealed values within the tolerable limits for uniformity, RC values in >2mm rods, and SORs in the non-radioactive water- and air-filled chambers. There was low variability in radiotracer uptake in all major organs of mice scanned using the four-animal bed versus those imaged using a single bed imaging platform.

**Conclusion:** Analysis of images acquired using the four-bed mouse hotel confirmed its utility to increase the throughput of small animal PET imaging without considerable loss of image quality and quantitative precision. In comparison to a single mouse bed, the cost and time associated with each scan were substantially reduced.

## Introduction

As a non-invasive imaging tool, positron emission tomography (PET) is used in preclinical research across multidisciplinary areas of work for whole-body, dynamic examination of biochemical processes under normal and pathophysiological conditions (*1-4*). As an important translational tool, preclinical PET has enabled the development of a multitude of different radiotracers that are currently in clinical use. For example, the development of radiotracers targeting PSMA for human prostate cancer imaging, theranostic approaches, and the ability to track therapeutic cells *in vivo* all stem from their thorough evaluation in rodent models (*5-7*).

A considerable limitation of current small-animal PET-computed tomography (PET/CT) imaging is the low throughput of image acquisitions. Single animal imaging becomes particularly restrictive when radioactive isotopes with short half-lives, such as carbon-11 and fluorine-18, or complex dynamic imaging studies are employed. Subsequently, to design sufficiently-powered studies, high costs accumulate. For many research Groups, these high imaging costs present a barrier for wide-spread preclinical adoption of PET and may restrict the frequency of radioactive preparations available to others that have invested in this imaging modality. A potential solution to this economic problem is to scan multiple animals simultaneously. A number of commercially-available preclinical scanners possess adequate axial length and diameter to achieve multi-animal imaging, and as a result, many user-designed multi-animal holders have entered into routine use (*8-10*). Often, however, these beds lack the thorough characterization required for the production of reproducible and quantifiable PET data, with animal heating and monitoring capabilities frequently omitted from the bed design. It is well recognised that temperature and anaesthesia can greatly impact the biodistribution of injected radiotracers and so maintaining control of these variables is essential to maintain reproducibility across subjects and between sites (*11*). It is also important to consider the impact on image quality and quantitative accuracy when multiple animals are scanned in the same field of view. The presence of more than one concentrated source of radioactivity my negatively affect attenuation and the number of scatter events, with reductions in resolution and sensitivity resulting as subjects are placed away from the centre field of view (CFOV).

To examine and overcome the low throughput of conventional PET imaging, a four-bed mouse ‘hotel’ was developed and validated. The animal hotel was designed to deliver an even distribution of anaesthesia to each nose cone and maintain animals at 37 °C. Heat and temperature tests were performed and the National Electrical Manufacturers Association (NEMA) NU-4 2008 standard protocol for small animal PET scans was used to assess the image quality of phantom PET scans. We then investigated whether the mouse hotel negatively impacted the quality of *in vivo* dynamic PET images simultaneously acquired with four mice. Following injection of [^18^F]2-fluoro-2-deoxyglucose (FDG) into female balb/c mice, images were subsequently compared to those acquired using a single animal bed.

## Materials and methods

### Design and development of the four-bed mouse hotel

We designed the four-bed mouse hotel for use in a Mediso nanoScan PET/CT (**Fig. 1A**). The hotel was designed using high temperature-resistant plastic and consisted of four holding chambers with individual nose cones for anaesthesia delivery (**Fig. 1B**). The bed contained chambers to allow the flow of heated air to all four beds within the animal hotel, with each mouse place equidistant from the CFOV and designed to allow the simultaneous imaging of up to four mice within the same single field of view (**Fig. 1C**). The modular design allowed the removal of the top bed layer when only 1-2 mice were required for imaging.

**FIGURE 1.**
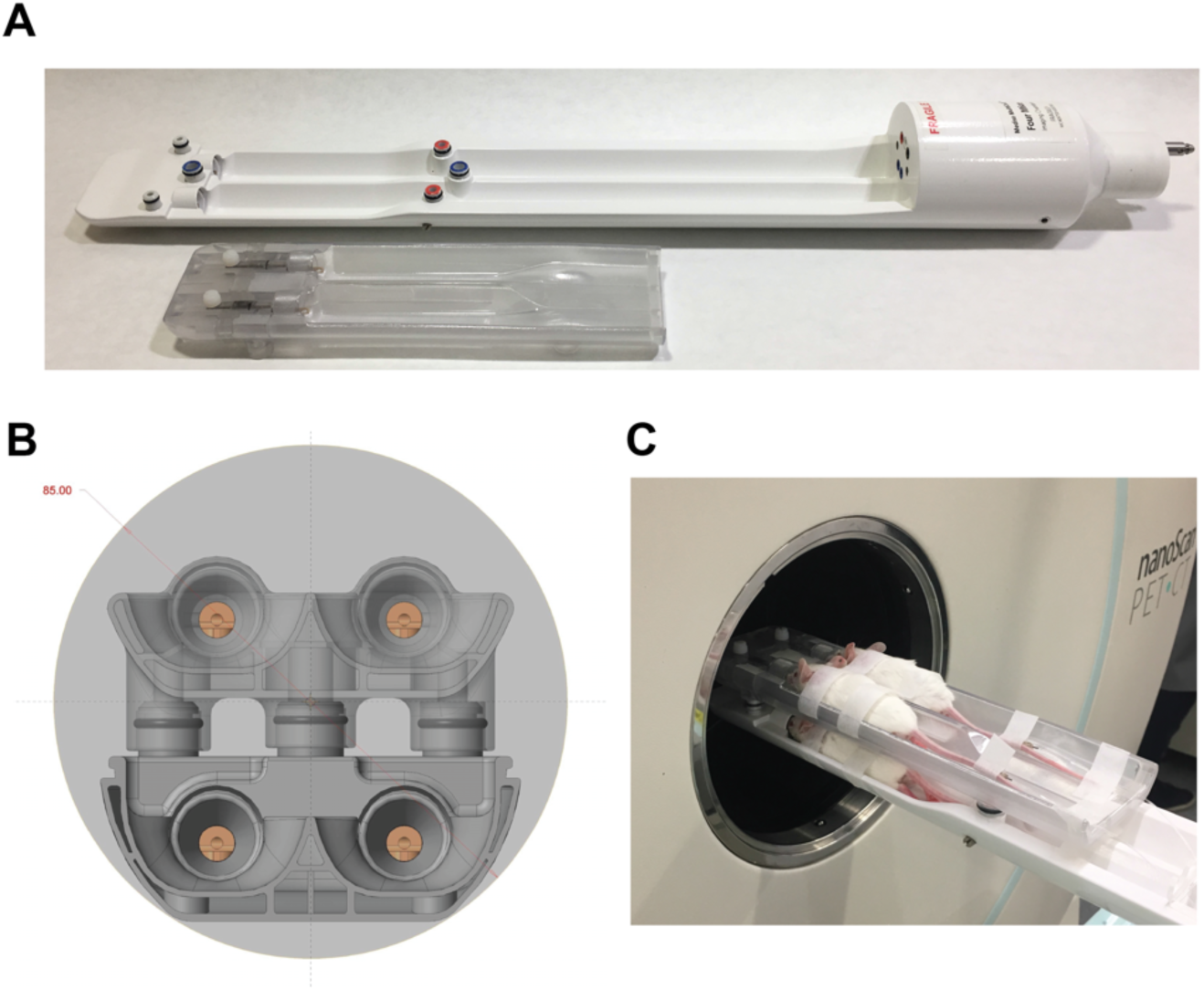
Design of the four-bed mouse hotel. (A) The modular design with removable top bed allowed between one and four 50 g mice to be simultaneously imaged. (B) Cross-section of the bed within the PET FOV (light grey). The four individual nose cones are visible, with the air-filled chamber located under each animal bed also in view. (C) Photograph of four female balb/c mice within the four-bed animal hotel.

### Temperature and anaesthesia tests

The air flow temperature delivered to the bed was set at 38°C and allowed to plateau for 5 min before temperature readings were taken using a thermal camera (FLIR systems AB-E60). Each animal holding bed within the hotel was measured individually at multiple positions. To measure anaesthesia delivery, a flowmeter (Dwyer RMA-26-68V) was attached to each individual anaesthesia nose cone.

### NEMA Mini Image Quality Phantoms studies

Cylindrical mouse-sized mini image quality phantoms were designed specifically for use in the four-bed mouse hotel by Mediso Medical Imaging Systems. Each phantom, made from Plexiglas, consisted of three parts: a large uniform compartment; a solid region with 5 fillable rods of 1 mm, 2 mm, 3 mm, 4 mm and 5 mm in diameter; and a non-radioactive region containing water- and air-filled chambers, designed to replicate the conditions of an *in vivo* PET/CT scan. Following the guidelines suggested by NEMA NU-4, each phantom was filled with 3.7 MBq of [^18^F]FDG in 10 mL of PBS (Thermo Scientific), decay-corrected to the start of acquisition. Clinical-grade [^18^F]FDG was obtained from PETNET solutions. All phantoms were thoroughly mixed and bubbles carefully removed before being placed onto the imaging bed. The phantoms were examined using different arrangements within the bed. For multiple phantom scans, four phantoms were placed in the animal hotel at the same time (configuration 1). For single phantom scans, one phantom was placed into each bed location within the hotel and imaged individually (configuration 2). Additionally, a single phantom was imaged using a single mouse bed for comparison (configuration 3).

### NEMA Mini Image Quality Phantoms studies

Four phantoms were imaged on the four-animal bed following the exact specification suggested by the PET manufacturer for single mouse imaging. Dynamic PET acquisition was performed on a Mediso nanoScan PET/CT over 20 min followed by CT (480 projections; 50kVp tube voltage; 300ms exposure time; 1:4 binning; helical acquisition). Whole body Tera-Tomo 3D reconstruction with 4 iterations and 6 subsets was performed (1-5 coincidence mode) using a voxel size of 0.3 mm^3^. Images were corrected for attenuation, scatter and decay. A gaussian filter of 0.7 was added to the post reconstructed PET images using VivoQuant software (v.2.5; Invicro Ltd.).

To optimise the manufacturer’s suggested scanner and reconstruction parameters, the tube voltage for CT imaging was increased to 70kVp. Additionally, whole body Tera-Tomo 3D reconstruction with 10 iterations and 6 subsets was performed (1-5 coincidence mode) using an isotropic voxel size of 0.3 mm^3^. Images were corrected for attenuation, scatter and decay. A gaussian filter of 0.7 was added to the post reconstructed PET images using VivoQuant software (v.2.5; Invicro Ltd.).

### NEMA NU-4 2008 tests

Post reconstruction, NEMA NU-4 2008 tests were performed to evaluate the effect of multiple subjects on scanner performance. Image noise was expressed as percentage standard deviation (%STD) by selecting a large FOV (75% of the active diameter) in the centre of the fillable region of the phantom. Activity recovery coefficients (RCs) in the five fillable rods were calculated from the maximum detected activity in each rod, divided by the mean total phantom activity concentration. To evaluate scatter correction, spill-over ratios in the non-radioactive water- (SOR_water_) and air-filled (SOR_air_) chambers were measured as the activity detected in these regions, divided by the mean total phantom activity concentration. Data were exported and analysed in Graphpad Prism (v.8.0).

### PET/CT animal imaging studies

All animal experiments were performed in accordance with the United Kingdom Home Office Animal (scientific procedures) Act 1986. PET acquisition was performed on a Mediso nanoScan PET/CT system. Female Balb/c mice aged 9-12 weeks (Charles River Laboratories) were fasted for 24 h prior to image acquisition. Mice were anaesthetized with 2.5% isoflurane in oxygen and a tail vein cannula inserted. A bolus of 3.7 MBq of [^18^F]FDG was injected in approximately 100 µL of PBS and dynamic images were acquired immediately over a period of 60 minutes. For attenuation correction and anatomical reference, CT images were acquired following PET imaging (480 projections; 70kVp tube voltage; 300 ms exposure time; 1:4 binning; helical acquisition;). Animals received 2.5% isoflurane in oxygen throughout the scan and were maintained at 37°C by the air flow heated bed. All animals were visually monitored throughout the imaging procedure.

The acquired data were sorted into 19 time frames of 4 × 15 seconds, 4 × 60 seconds, and 11 × 300 seconds for image reconstruction (Tera-Tomo 3D reconstructed algorithm; 4 iterations; 6 subsets; 400-600 keV; 0.4 mm^3^ voxel size). VivoQuant software (v.2.5, Invicro Ltd.) was used to analyse reconstructed images. Regions of interest (ROIs) were drawn manually using CT images and 30-60 minute summed dynamic PET images as reference. Time versus radioactivity curves (TACs) were generated using normalised count densities to the injected activity and the area under the time versus radioactivity curve (AUC) generated.

### Statistics

All data were expressed as the mean ± one standard deviation (SD). Statistical significance was determined using either a two-tailed t-test or analysis of variance (ANOVA) followed by t-tests multiple comparison correction (Tukey method; GraphPad Prism v.8.0).

## Results

### Physiological regulation

The animal bed was designed to circulate heated air evenly across the four animal chambers. This design facilitated a constant and even temperature distribution to all four animal holding beds (36.8 ± 0.4 °C; *n* = 4; **Fig. 2A**). As a design consideration, air inlets, where temperatures over 37 °C were observed, were placed away from the region of the bed where the mice were positioned. Flow meter readings showed anaesthesia was distributed evenly to each nose cone (2.8 ± 0.1 L/min; from the four nose cones; **Fig. 2B**).

**FIGURE 2.**
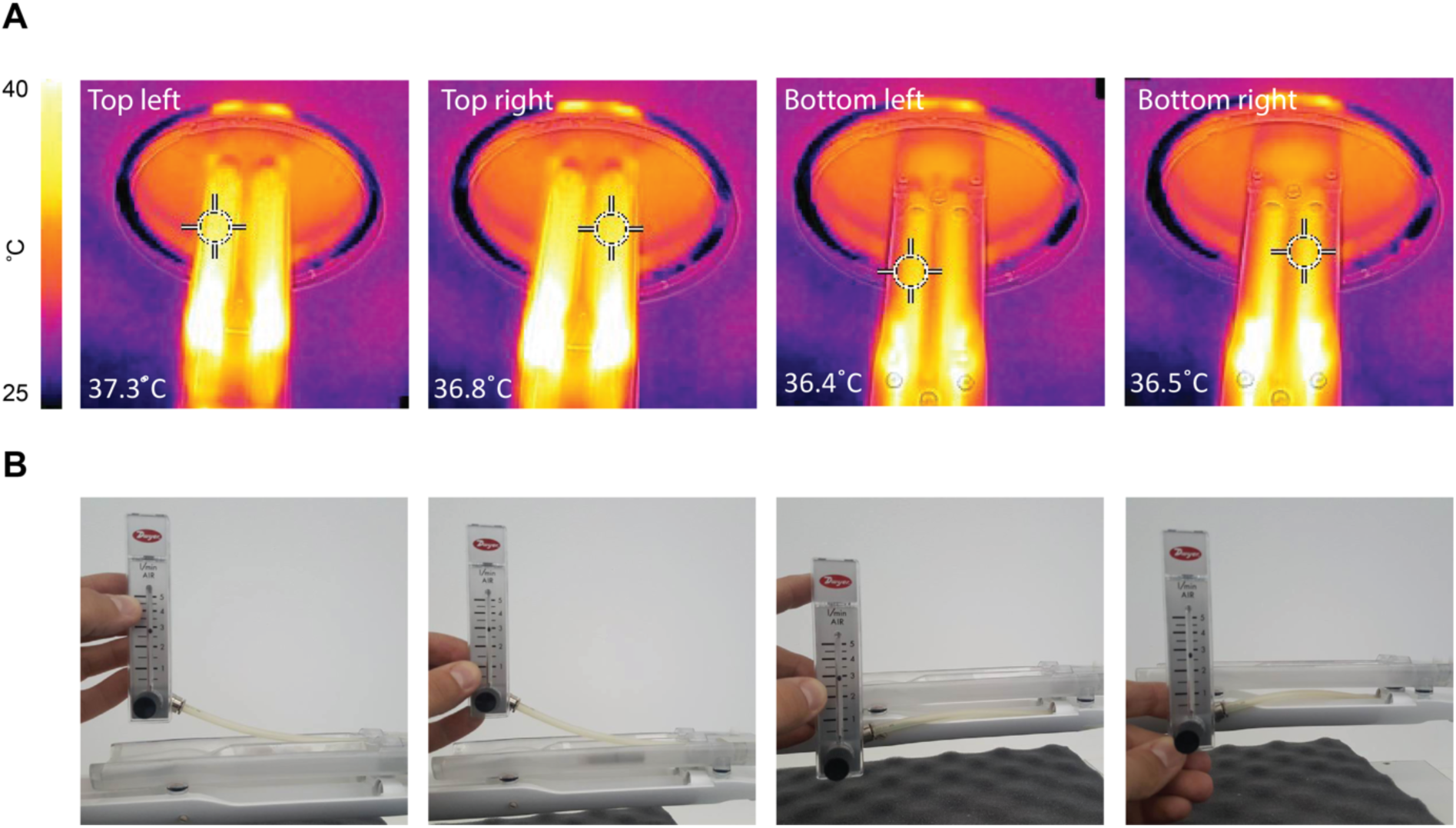
Temperature and anaesthesia flow rate uniformity testing in the four-animal bed. (A) Temperature measurements were acquired using a thermal camera following heating of the four-bed mouse hotel. The bed temperature for the bottom bed layer was assessed following removal of the top layer. Elevated temperature was evident by the air inlets that were positioned away from the mice towards the back of the bed. (B) Flowmeter values representing anaesthesia output from each nose cone.

### Phantom studies

The bed’s performance was evaluated using mini image quality NEMA NU-4 phantoms specifically designed for their use in the four-bed mouse hotel (**Fig. 3A**). Each phantom contained three discrete regions, holding a total volume of 10 mL. Quantitative analysis was performed using NEMA NU-4 tests following reconstruction, with the tolerable limits set by NEMA shown in **Supplementary Table 1**. Initially, phantom images were reconstructed using those parameters recommended by the manufacturer for standard scans on a single animal bed (**Supplementary Methods**). The NEMA NU-4 test results using these standard parameters are shown in **Supplementary Figure 1**. Subsequently, these reconstruction parameters were optimised to improve image quality by increasing the number of iterations and subsets to 10 and 6, respectively. The RC values measured in the five fillable rods with diameters of 1-5 mm were used to evaluate spatial resolution of images acquired using the animal hotel and were compared to a single animal bed. Reduced spatial resolution was evident in phantoms imaged using the four-bed mouse hotel when examining the RCs of the 1 mm and 2 mm rods, with values falling outside the tolerable limits suggested by the NEMA NU-4 guidelines (0.06 ± 0.01 and 0.68 ± 0.07, respectively; *n* = 4). Spatial resolution, however, was not affected in the larger rods (1.08 ± 0.09, 1.06 ± 0.05, 1.05 ± 0.04, for 3 mm, 4 mm and 5 mm rod, respectively; *n* = 4), with values similar to those acquired using a single mouse bed (1.05 ± 0.11, 1.04 ± 0.05, 1.02 ± 0.04; *n* = 4; **Fig. 3B**). Interestingly, when imaging a single phantom in the four bed mouse hotel, with the exception of the 3 mm rod, the RC values were all within the tolerable limits at all rod diameters tested, presumably due to the presence of a single point-source of radioactivity (0.23 ± 0.02, 0.99 ± 0.09, 1.2 ± 0.08, 1.1 ± 0.08 and 1.1 ± 0.03 for 1 mm, 2 mm, 3 mm, 4 mm and 5 mm rods, respectively; *n* = 4).

**FIGURE 3.**
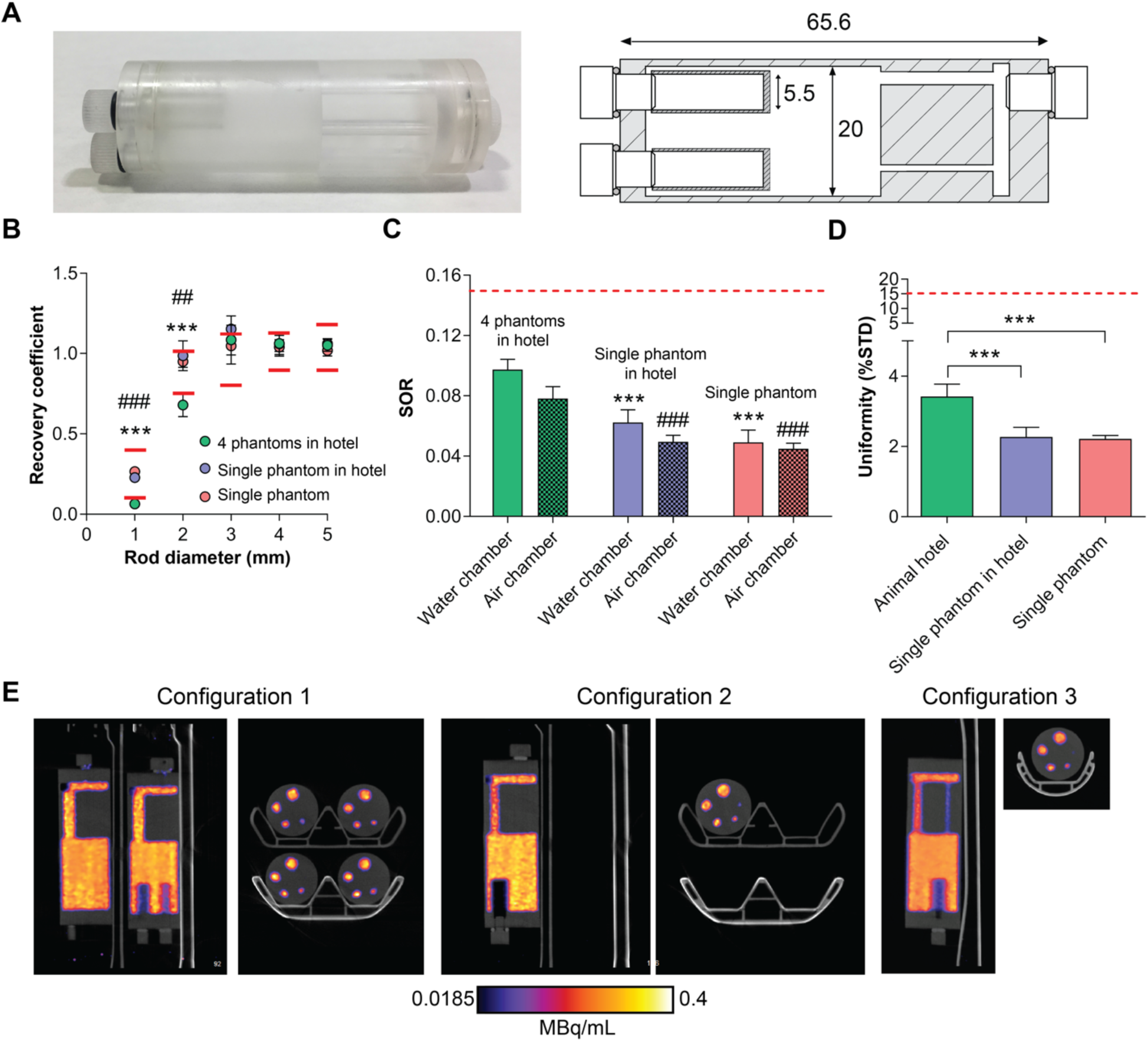
Evaluation of image quality using mini IQ NEMA NU-4 phantoms. Mini IQ phantoms were designed specifically for the evaluation of the four-bed mouse hotel (A). Dimensions are shown in mm. Mini IQ phantoms were filled with ~3.7MBq of [^18^F]FDG and imaged over 20 mins. Following reconstruction, NEMA NU-4 tests were performed. The RC values of 1-5 mm rods (B), SORs (C) and uniformity values (D) of phantoms imaged using the four-bed mouse hotel and a single mouse bed were acquired and compared. (E). Representative sagittal and axial PET images of 0-20 min summed activity of imaging configurations 1, 2 and 3. Red lines represent the tolerable limits set by NEMA NU-4. For (B), *** = *P* < 0.001 for configuration 1 *vs* configuration 2; ### = *P* < 0.001 for configuration 1 *vs* configuration 3; for (C), *** = *P* < 0.001 for configuration 1 SOR_water_ vs configuration 2 and 3; ### = *P* < 0.001 for configuration 1 SOR_air_ vs configuration 2 and 3.

The SORs for both the water- and air-filled chambers were within the tolerable limit of 15 % for all imaging configurations, suggesting an appropriate scatter correction was applied. SORs for the four-bed mouse hotel were significantly higher than those acquired using a single mouse bed, however. For configuration 1, 2 and 3, the SOR_water_ was 9.7% ± 0.7%, 6.2% ± 0.8%, 4.9% ± 0.8%, respectively (*P* = 0.0003 and *P* < 0.0001, for configuration 1 vs. configuration 2 and configuration 1 vs. configuration 3, respectively). The SOR_air_ was 7.8% ± 0.08%, 4.9% ± 0.4%, 4.5% ± 0.4%, for configuration 1, 2 and 3, respectively (*P* = 0.0001 and *P* < 0.0001, for configuration 1 vs. configuration 2 and configuration 1 vs. configuration 3, respectively; *n* = 4; **Fig. 3C**). The variation in activity concentration, as measured by the % STD in the uniform region of the phantom, was higher when four phantoms were imaged simultaneously compared to the single phantom configurations, representing elevated image noise (3.4% ± 0.35%, 2.3% ± 0.27% and 2.2% ± 0.1% for configuration 1, 2 and 3, respectively; *P* = 0.003 for configuration 1 vs. 2; *P* = 0.004 for configuration 1 vs. 3; *n* = 4; **Fig. 3D**). All values, however, were well below the tolerable limit of 15%. Representative single-slice [^18^F]FDG PET/CT images (0 – 20 min summed activity) of the mini IQ phantoms are displayed in **Fig. 3E**, comparing the three imaging configurations.

### Animal studies

To evaluate the bed under standard *in vivo* imaging conditions, four healthy mice were simultaneously imaged with [^18^F]FDG PET dynamically over 60 min in the four-bed mouse hotel (**Fig. 4A**). An additional four mice were imaged in a single mouse bed for comparison (**Fig. 4B**). Following reconstruction, TACs revealed low levels of variability in [^18^F]FDG uptake for all major organs for animals imaged in the four-bed animal hotel (**Fig. 4C**). There was no significant difference in the area under the TAC for all major organs with the exception of muscle uptake (87.3 ± 36.3 vs. 34.1 ± 17.1 %ID·h/g; *P* = 0.04; *n* = 4; **Supplementary Fig. S2**). When compared to the single animal scans, the variation in radiotracer tissue uptake between subjects increased with the mouse hotel. Representative maximum intensity projections following the manual removal of the bed are shown in **Supplementary Fig. S3**.

**FIGURE 4.**
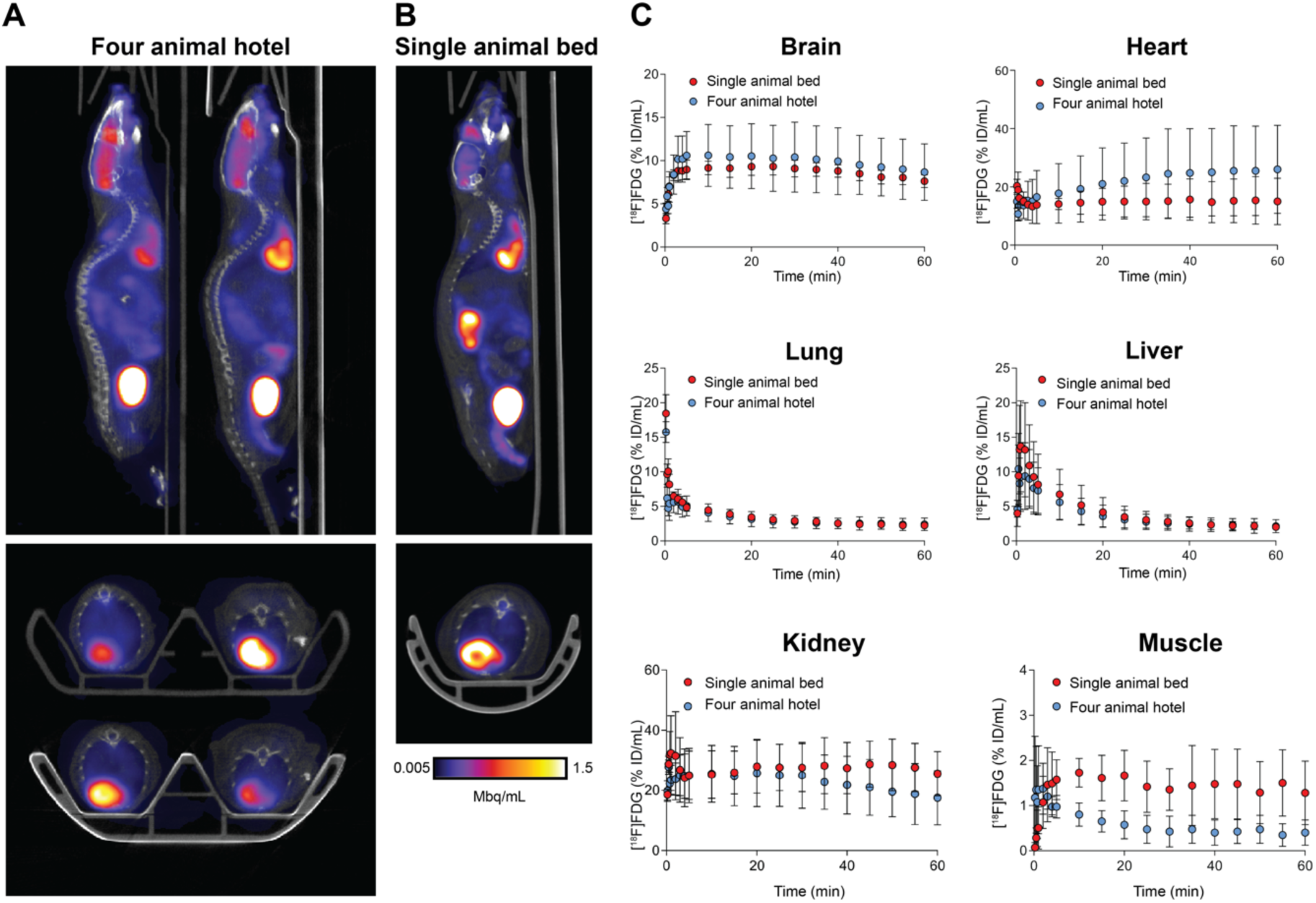
*In vivo* validation of the four-bed mouse hotel and comparison to a single mouse holder. Representative sagittal and axial PET images of 30-60 min post injection of mice imaged using (A) the four-bed mouse hotel and (B) a single mouse bed. Br, brain; H, heart; K, kidney; B, bladder. (C). Time *vs.* radioactivity curves (TACs) of major organs of interest normalised to the percentage injected activity. Error bars represent one s.d. from the mean value (*n* = 4 animals).

## Discussion

Here, we sought to validate the performance of a commercially-designed four-bed mouse hotel for use in a Mediso nanoScan PET/CT imaging system. Utilizing a four-bed mouse hotel for PET/CT small animal imaging has a major benefit over conventional single animal imaging, saving investigators’ time and reducing overall costs. Additionally, by increasing the number of animals that can be scanned for the same cost and time, the statistical power and corresponding confidence in the imaging data can be substantially increased.

The feasibility of imaging multiple animals in the same small animal PET/CT imaging system has been reported in multiple studies (*8-10,12*), facilitated through the development of new small animal PET/CT imaging platforms which offer sensitive and reproducible imaging over both a large axial and transaxial FOV. These large axial FOVs enable several animals to be imaged simultaneously. To the best of our knowledge, however, physiological regulation and monitoring has not been incorporated into the user-developed animal holders currently in use. Unlike previous unheated animal bed holders which can only be used for a short duration (*12*), dynamic imaging over multiple hours is achievable on this four bed mouse hotel as a result of fine temperature regulation. Changes in physiological conditions can affect mammalian physiology, including disruption to thermoregulation, respiration and cardiac output (*13*). Maintaining a constant bed temperature will therefore limit variations in radiotracer pharmacokinetics, whilst minimizing any potential pain, suffering, distress or lasting harm that may be caused to the animal.

To evaluate the performance of the bed, phantom tests were performed under the conditions of the NEMA NU-4 2008 standard protocol for small-animal PET systems. The NEMA NU-4 test results acquired using the suggested reconstruction parameters set by the NEMA guidelines are provided in **Supplementary Fig. S1**. To improve upon the data acquired using the manufacturer’s suggested parameters for a conventional scan using the single-mouse bed, the reconstruction parameters were altered, with the number of iterations increased from four to ten. Following completion of the NEMA tests on the newly reconstructed images, low-level radioactivity was detected in the water and air chambers within the NEMA phantoms due to the scattering of photons. The SOR in both the air and water chambers was increased in phantoms imaged using the four-bed mouse hotel compared to the single-bed images, which can be attributed to the increased number of subjects in the FOV, leading to elevated photon scattering. The amount of scatter detected in the non-radioactive chambers of phantoms imaging using the four-bed mouse hotel, however, was below the maximum tolerable limit, suggesting minimal loss in image quality.

To assess the expected impact on spatial resolution in the mouse hotel, RCs values for the 1-5 mm rods in the mini IQ phantom were calculated. For the 1 mm and 2 mm rods, the RC values were decreased by 76% and 26%, respectively, compared to phantoms scanned on a single mouse bed. For the larger diameter rods, all values fell within the tolerable limits. A reduction in spatial resolution is expected with the four-mouse bed as each imaging chamber is placed away from the CFOV. Matching previous findings (*14*), increasing the number of iterations from four to ten during reconstruction improved the RCs of all rods, however, this amplified image noise and subsequently reduced observable image quality. During small animal PET imaging of tumor-bearing mice, lowered spatial resolution is unlikely to alter quantitative information as most tumors imaged are typically larger than 50 mm^3^; increasing the number of iterations during reconstruction therefore may not be beneficial. Care, however, should be taken when evaluating radiotracer uptake in small tissues, such as lymph nodes or small metastatic lesions when using the four-animal bed. Here, [^18^F]FDG radiotracer uptake in the major organs of healthy mice scanned simultaneously versus those imaged individually produced similar quantitative values, despite the expected biological variation in both radiotracer retention and excretion. To ensure maximum image quality, applying scatter and attenuation correction to all *in vivo* imaging studies using the four-bed mouse hotel is highly recommended. Unlike in the phantom studies, image reconstruction was not optimised for the *in vivo* imaging studies performed, with refinement of reconstruction algorithms likely to further improve image quality.

Whilst we have demonstrated the feasibility of simultaneous imaging of up to four mice whilst maintaining quantitative precision, there are a number of improvements in bed design that should be incorporated prior to commercialization. At the time of data acquisition, both breathing and heart rate monitoring was available for only one of the four mice using this prototype bed. An updated bed that incorporates physiological monitoring for all four mice has subsequently been developed. An additional consideration is the potential degradation of image quality when four animals are imaged simultaneously. Here, phantom imaging studies revealed potential issues with image reconstruction, leading to loss of data quality. Suboptimal image reconstruction and lowered image resolution was initially seen through the loss of detected radioactivity in the small 1 mm and 2 mm rods of phantoms scanned in the four-bed mouse hotel. Furthermore, in our *in vivo* studies, reduced sensitivity became apparent when comparing the muscle TACs of animals scanned using the four bed mouse hotels versus a single mouse imaging bed. The lowered radioactivity detected in the muscles of mice scanned in the mouse hotel when compared to those scanned on the single bed can be attributed to exaggerated partial volume effects from small ROIs containing low concentrations of radiotracer.

Despite some minor drawbacks, a considerable benefit of using the four-bed mouse hotel during PET imaging are the large cost savings that can be made compared to a conventional single animal scanner. A clinical dose of ~370 MBq [^18^F]FDG on average costs ~£300 in the UK. Considering the time required to position the animal on the bed, time taken to prepare the radioactive dose radioactive dose and a typical 10 min CT scan, with a constantly-decaying radiotracer, typically each clinical dose will allow for approximately five 60 min dynamic scans. For novel radiotracer development and discovery PET imaging where the radioactivity concentration received may be substantially less than a clinical dose of [^18^F]FDG, the number of consecutive scans that can be achieved is even fewer. For sufficiently powered studies, a sample group of >eight are typically required to identify moderate changes in radioactive concentration with statistical confidence. Performing 60-minute dynamic [^18^F]FDG PET/CT on a conventional single mouse bed would require a minimum of two radioactive preparations to achieve this statistical power. Additionally, imaging institutes may charge ~£200 per hour for use of the scanner. To image eight mice using a single animal bed, a total of 12 hours split over two days would be required. However, imaging eight mice using the four-bed mouse hotel would only require a maximum of four hours in one day. In total, by using the four-bed mouse hotel, we estimate scanning/radiotracer costs could be reduced by approximately 60% using commercially-available [^18^F]FDG, with far higher savings potentially achieved with low-yielding novel radiotracers.

## Conclusion

A four-bed mouse hotel was designed to aid imaging scientists conduct their research in a more time-efficient, cost-effective manner by quadrupling the number of mice that can be imaged in a single session. In particular, when performing experiments using short-lived isotopes, multiple animals can be scanned using a single synthesis, which would otherwise not be possible. The design of the four-bed mouse hotel allowed for uniform control over temperature and anaesthesia, with phantoms studies and *in vivo* imaging of mice confirming its utility to increase the throughput of small animal PET imaging without considerable loss of image quality and quantitative precision. In comparison to a single mouse bed, cost and time associated with each scan were substantially reduced.

## Disclosure

Authors Gábor Mócsai, Zoltán Nyitrai and Sandor Hobor are employed by Mediso Medical Imaging Ltd, but do not own any shares or have any additional financial benefits in addition to general compensation for employment. They do not hold any patents related to the topic of the publication. None of the other authors have a conflict of interest to declare. This study was funded through a Wellcome Trust and Royal Society Sir Henry Dale Fellowship (107610/Z/15/Z) and a CRUK UCL Centre Non-Clinical Training Award (A23233) to Timothy H. Witney.

### Acknowledgements

The authors would like to thank Mike Bewick, Max Capel and Gabor Nemeth for insightful scientific discussions and Tammy Kelber for help with *in vivo* experiments.

