## Supplemental data for "High Throughput PET/CT Imaging Using a Multiple Mouse Imaging System"

Supplemental table 1. Tolerable NEMA NU-4 values as set by NEMA for image quality phantoms.

| Specifications | | |
|---|---|---|
| Uniformity (%STD) | | 15 |
| Spill over ratio | Air filled | <0.15 |
| | Water filled | <0.15 |
| Recovery coefficients | 1 mm | 0.1-0.4 |
| | 2 mm | 0.75-1.0 |
| | 3 mm | 0.8-1.1 |
| | 4 mm | 0.9-1.15 |
| | 5 mm | 0.9-1.19 |

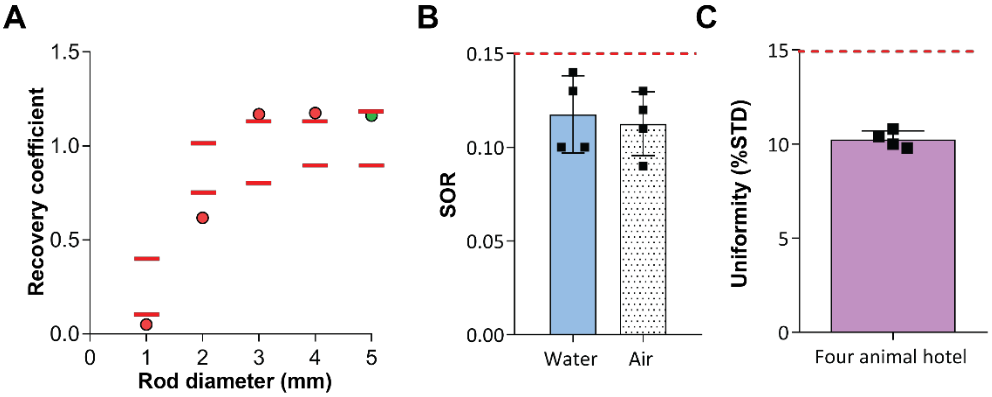


**Figure S1. NEMA NU-4 tests of phantom images following reconstruction using manufacturer-suggested parameters.** Phantoms were imaged with [$^{18}$F]FDG PET over 20 min in the four-bed mouse hotel, followed by CT (480 projections; 50kVp tube voltage; 300ms exposure time; 1:4 binning; helical acquisition). Phantom images were reconstructed using parameters suggested by the manufacturer for standard acquisitions (whole-body Tera-Tomo 3D reconstruction, 4 iterations and 6 subsets was performed, 1-5 coincidence mode). NEMA NU-4 tests were performed on the reconstructed images. **A**. The RCs of 1-4 mm rods fell outside of the tolerable limits, with only the 5 mm value passing the NEMA NU-4 test. For both $SOR_{water}$ and $SOR_{air}$ values (**B**) and uniformity (**C**) values were below the tolerable limits (red dashed lines).

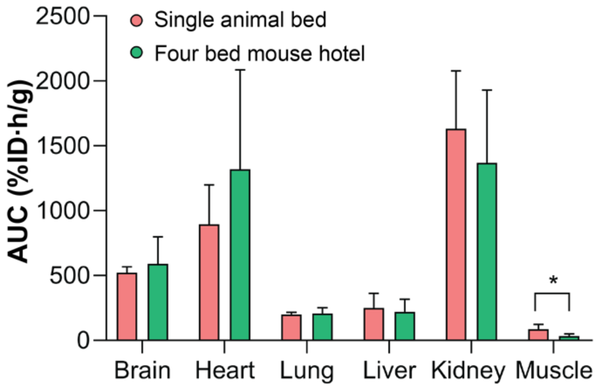


**Figure S2. Time activity area under the curve values for organs with substantial levels of [$^{18}$F]FDG uptake.** Data expressed as the mean plus standard deviation. *n* = 4 animals per group.

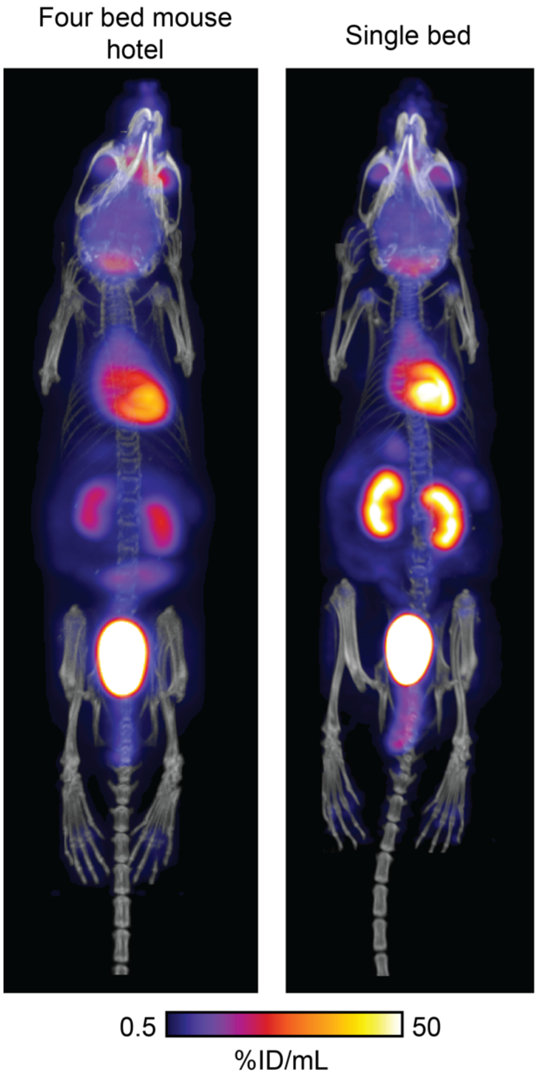


**Figure S3**. Representative [$^{18}$F]FDG PET/CT maximum intensity projection 30-60 min post injection of a healthy balb/c mouse imaged using the four-bed mouse hotel and a single imaging bed. The bed has been manually cropped out of both images.
